# Zonal Patterning of Extracellular Matrix and Stromal Cell Populations Along a Perfusable Cellular Microchannel

**DOI:** 10.1101/2024.07.09.602744

**Authors:** Brea Chernokal, Bryan J. Ferrick, Jason P. Gleghorn

## Abstract

The spatial organization of biophysical and biochemical cues in the extracellular matrix (ECM) in concert with reciprocal cell-cell signaling is vital to tissue patterning during development. However, elucidating the role an individual microenvironmental factor plays using existing *in vivo* models is difficult due to their inherent complexity. In this work, we have developed a microphysiological system to spatially pattern the biochemical, biophysical, and stromal cell composition of the ECM along an epithelialized 3D microchannel. This technique is adaptable to multiple hydrogel compositions and scalable to the number of zones patterned. We confirmed that the methodology to create distinct zones resulted in a continuous, annealed hydrogel with regional interfaces that did not hinder the transport of soluble molecules. Further, the interface between hydrogel regions did not disrupt microchannel structure, epithelial lumen formation, or media perfusion through an acellular or cellularized microchannel. Finally, we demonstrated spatially patterned tubulogenic sprouting of a continuous epithelial tube into the surrounding hydrogel confined to local regions with stromal cell populations, illustrating spatial control of cell-cell interactions and signaling gradients. This easy-to-use system has wide utility for modeling three-dimensional epithelial and endothelial tissue interactions with heterogeneous hydrogel compositions and/or stromal cell populations to investigate their mechanistic roles during development, homeostasis, or disease.

## Introduction

Developmental patterning of organs is orchestrated by biochemical and biophysical cues within and between tissues. Spatial gradients of soluble signaling molecules, called morphogens, are widely recognized to regulate gene expression, influencing cell differentiation and organ morphogenesis, such as budding and branching in tubular organs including the lung and kidney (1–3). Morphogen gradients develop via regional variations in the microenvironmental niche, including stromal cell populations and extracellular matrix (ECM) properties, which impart spatially distinct reciprocal cell-cell and cell-ECM signaling and interactions. These gradients and regional differences thereby direct patterning and differentiation events within the tissue (4). This is evident in kidney development, where β-catenin signaling activity is responsible for both nephron progenitor cell maintenance and differentiation depending on the concentration of the Wnt9b ligands secreted from the neighboring epithelium and interstitial cells (5–7). These gradients are further reinforced by the formation of 17 unique mesenchymal cell populations arranged in zones along the developing nephron that differentially express ECM-modifying enzymes (8) to pattern ECM stiffness along the cortical-medullary axis (9). Whereas specific developmental signaling pathways have been elucidated, their coordination and relationship to physical cues within the local microenvironment are poorly understood.

Our knowledge of the complex and integrated mechanisms that underpin development are largely derived from investigations of model organisms, including mice, *C. elegans*, and *Drosophila*, and more recently, human organoid systems. The ability to genetically manipulate these systems allows for system-level understanding. However, their inherent complexity and confounding variables have rendered the dissection of the contribution of individual microenvironmental niches challenging. *In vitro* models created with morphogen or mechanical gradients within a bulk hydrogel traditionally used to investigate their role in development or disease are often 2D, which lack architectural cues relevant to many tissues, including the kidney. Microphysiological systems (MPSs), while reductionist, offer a platform to recapitulate and precisely control key biochemical, biophysical, and cellular components of the microenvironment with well defined geometries (10–12). However, current techniques to regionally pattern the ECM around a microchannel within an MPS only have control radially/circumferentially and not longitudinally along the microchannel length. This perpendicular patterning does not reflect the morphogen gradients, ECM composition, or mesenchymal cell population patterning observed along the developing epithelial tube. For instance, the cortical-medullary axis of the developing epithelial nephron has distinct niches across lengthscales of microns to millimeters. To interrogate the role of these patterned niches on epithelial development and morphogenesis, it is necessary to recapitulate these features and their length scales. Therefore, an MPS model capable of recapitulating the zonal patterning of the surrounding ECM properties and cell identity along the longitudinal axis of an epithelialized microchannel is needed to understand the relationship between the developmental signaling pathways and these local microenvironment cues.

We have thus created and validated a MPS with the ability to spatially pattern ECM and stromal cell populations along the length of a continuous and perfusable micron-sized epithelial tube. We fabricated a polydimethylsiloxane (PDMS)glass microfluidic device, by utilizing a 3D mold that defines the hydrogel region including a series of orthogonal gel in-jection ports, which enabled serial patterning of multiple hydrogels with unique composition and/or encapsulated cellular populations along the microchannel length. Validation of the device using both cellular and acellular gels, confirmed that the gel-gel interfaces did not impede morphogen diffusion, microchannel lumenization, or media perfusion. Finally, we validated the use of the device by demonstrating regional epithelial phenotypic changes in response to a spatially patterned stromal cell population. Specifically, epithelial sprouting into the hydrogel stroma from the continuous epithelial tube was regionally induced only within discrete zones of spatially patterned stromal fibroblasts. The results of this study confirm that this MPS is a viable model system for constructing and interrogating the role of stromal cell and ECM identity, composition, and biophysical properties in development, homeostasis, and disease.

## Materials and Methods

### Device fabrication

Device molds were designed in Solidworks 2021 (Dassault Systèmes) then 3D printed with TR250LV high-temp resin (Phrozen Technology) using an 8k resolution masked stereolithography printer (Phrozen Technology). The mold forms were printed parallel to the build plate to minimize transfer of layer line artifacts and increase optical clarity when replica molded in polydimethylsiloxane (PDMS, Sylgard 184; Dow Corning). Molds were washed with isopropyl alcohol for 10 minutes to dissolve any uncured, liquid resin on the parts. While the molds were still soft, two 400 µm holes were manually drilled into the left and right sides at the cross-section’s centroid for later placement of a stainless-steel 400 µm OD hypodermic tube (McMaster-Carr). Molds were then cured under UV light for 20 minutes at 60°C, followed by an additional 24 hours of curing at 50°C to prevent resin leachate, which would inhibit PDMS curing during casting (13). Following curing, a 2-3 cm length of the 400 µm OD hypodermic tube was placed into each drilled port to create microchannel needle guides within the PDMS. The 3D-printed mold was cast in PDMS at a 1:10 curing agent to prepolymer ratio, then degassed in a vacuum chamber for 1 hour and cured at 60°C overnight. Biopsy punches were used to create inlets and outlets for the gel injection ports (1.5 mm), air relief ports (1.5 mm), and media wells (7 mm). The PDMS devices were plasma bonded to glass coverslips following standard techniques (14). After bonding, devices were cleaned with 70% ethanol for 30 minutes, rinsed with sterile phosphate buffered saline (PBS; Corning), then placed under UV for 30 minutes to ensure sterility. Device molds consisted of either small devices (7.5 mm long channel) with two hydrogel injection ports and large devices (15 mm channel) with two or four injection ports to produce two or four zones along the length of the epithelial tube.

To generate devices with a central hydrogel microchannel, a 140 or 160 µm acupuncture needle was treated with 1% bovine serum albumin (BSA; Fisher Scientific) for 30 minutes on ice to passivate the needle. An acupuncture needle was inserted into the device via the guide channels made from the hypodermic tubing. Devices for use with encapsulated cells in the bulk hydrogel were incubated with 2% polyethyleneimine (PEI; Sigma Aldrich) for 30 minutes, rinsed 3x with sterile water (HyClone Water; Cytiva), then incubated with 0.5% glutaraldehyde (GA; Fisher Scientific) for 45 minutes and rinsed with sterile water 3x before hydrogel injection to prevent delamination of the hydrogel from the PDMS due to cell-mediated contractile forces (15). Devices without cells encapsulated in the hydrogel bulk did not require any conjugation chemistry to bond the hydrogel to the PDMS sidewalls.

### Hydrogel formulation and zonal patterning within devices

Collagen hydrogels were made at either 3 mg/mL or 6 mg/mL density from rat tail collagen type 1, as previously described (14). Briefly, rat tail tendons were dissected and washed with PBS and isopropyl alcohol before being dissolved in acetic acid. Dissolved collagen was then lyophilized and stored at −80°C until use. A stock collagen solution was created by dissolving the lyophilized collagen in 0.1% acetic acid at a concentration of 8 mg/mL. On ice, collagen was diluted to the proper concentration in 10X Hanks Buffered Saline Solution (HBSS; ThermoFisher) and Dulbecco’s Modified Eagles Medium with 4.5 g/L glucose, sodium pyruvate, and Lglutamine (DMEM; Cytiva) and neutralized to 7.5 pH using 1 N NaOH. Fibrin hydrogels were made by mixing 10 mg/mL stock fibrinogen (Sigma Aldrich) 1:1 with PBS to create a 5 mg/mL fibrinogen solution. 5 mg/mL fibrinogen was then mixed with thrombin (50 U/mL; Cayman Chemical) at a 50:1 ratio (fibrinogen:thrombin).

For hydrogel addition into devices, all steps were performed in a biosafety cabinet. When using collagen hydrogels, the devices, reagents, and supplies were kept on ice to increase to the working time of the collagen prepolymer and help ensure uniform polymerization. To zonally pattern two-region devices, two 1 mL syringes with blunt tip 18-gauge Luer-Lock needles (McMaster-Carr) were loaded with the mixed hydrogel solution, and each solution was added serially to the device. The first hydrogel was injected via one injection port until it reached the edge of the air relief channel. The second hydrogel was injected via the other injection port until it coalesced with the first and filled the air relief channel. Patterning of four region devices was achieved similarly, with hydrogels injected serially from left to right across the device. Following hydrogel addition, devices were immediately placed in the incubator at 37°C for 30 minutes. Following hydrogel polymerization, acupuncture needles were carefully withdrawn, creating a microchannel where cells could be seeded. The media reservoirs were filled with either DMEM culture media for cellular devices or PBS for acellular devices to prevent the hydrogel desiccation.

### Cellular seeding of devices

All cellular devices were made using small or large two-region devices. For co-cultured devices with cell-patterned stromal cell regions, NIH3T3 fibroblasts (3T3, ATCC); transduced to express green fluorescent protein (GFP, Addgene: 72263) were cultured in DMEM supplemented with 10% fetal bovine serum (FBS) and 1% penicillin and streptomycin at 37°C and 5% CO_2_. 3T3 cell suspension was added to the collagen prepolymer solution in place of the DMEM to achieve a final cell density of 5*10^5^ cells/mL. The solution was mixed carefully to suspend 3T3s throughout the collagen homogenously. It was then injected via one port of the device, and an acellular collagen hydrogel was injected on the other port, as described above. Following hydrogel polymerization, microchannels were seeded with Madin-Darby Canine Kidney (MDCK; ATCC) epithelial cells or MDCK cells transduced to express mCherry fluorescent protein (MDCK^mCherry^, Addgene: 72264). Both MDCK types were cultured in DMEM supplemented with 10% fetal bovine serum (FBS) and 1% penicillin and streptomycin at 37ºC and 5% CO_2_. Prior to removal of the acupuncture needle from the device, 100 µl of 10*10^6^ cells/mL MDCK cell suspension was added to one media reservoir and 80 µl of media to the second media reservoir. Once the acupuncture needle was removed, the volume difference established a hydrostatic pressure gradient to induce the flow of the cells at a controlled rate for seeding the channel. Devices were incubated for 5-10 min at 37°C to allow MDCKs to settle and adhere within the channel. The cells in each media well were then resuspended and the pressure gradient was reestablished before the devices were flipped to coat the top of the microchannel with MDCKs. Devices were again incubated for 5-10 min at 37°C. Following seeding of the microchannel, excess cell suspension was removed from both wells and replaced with fresh media. Devices were cultured under static conditions until full microchannel epithelialization, approximately five days with daily media changes, before being cultured under perfusion via a peristaltic pump (Ismatec). For pump perfusion, 1/16 in ID silicone tubing (McMaster-Carr) was connected to a 24-gauge blunt tip Luer-Lock needle and primed with culture medium before being inserted into one side of the device through the microchannel guide port within the PDMS. Medium at the outlet was passively collected. Devices were perfused at a constant flow rate of 5 µL/min for 24 hours. 3T3-MDCK coculture devices were cultured under static conditions with daily media changes.

### Epithelial tube and sprout quantification

Following seven days of static culture, live differential interference contrast (DIC) brightfield images of 3T3-MDCK co-culture devices were captured using a 20x/0.8 objective on a Zeiss Axio Observer Z1 widefield microscope. From these images, the quantity and locations of tubulogenic sprouts of the MDCK epithelium were recorded within 5 mm on either side of the gel-gel interface to compare the 3T3 seeded and unseeded regions. Images were analyzed in Zeiss ZEN software, and tubulogenic sprouts were defined as MDCK protrusions of at least 20 µm in length from the basal edge of the epithelial microchannel into the surrounding hydrogel. Microchannel diameters were measured in ZEN every 1 mm along the 15 mm length of each device from tiled images taken with a 5x/0.16 objective on a Zeiss Axio Observer Z1 widefield microscope.

### Immunofluorescent antibody staining

Devices were carefully rinsed three times with 1X PBS with calcium and magnesium (PBS Ca^2+^/Mg^2+^; Fisher). After washing, devices were fixed with 4% paraformaldehyde (PFA, ThermoFisher) with 0.1% Triton-X 100 (Sigma-Aldrich) at room temperature for 30 minutes. Following fixation, the devices were blocked and stained with anti-E-Cadherin (DECMA-1, Santa Cruz) at 4 °C overnight. Devices were again washed 3x with 1X PBS Ca^2+^/Mg^2+^; then secondary antibody (Anti-IgG polyclonal Rat DyLight® 650; ImmunoReagents) was added to the devices and incubated overnight at 4 °C. DAPI (ThermoFisher) and phalloidin (DyLight 488/555; Cell Signaling) were then incubated at room temperature for 1 hour. Devices were kept hydrated in PBS with Ca^2+^/Mg^2+^ at 4°C until imaging. Zstack images were taken on a Zeiss LSM800 confocal microscope with a 20x/0.8 objective.

### Interstitial flow assay

Small devices were made with one (single) or two patterned (dual) collagen (3 mg/mL) or fibrin (5 mg/mL) gels. 100 µl of 12.5 µg/mL 70 kDa Texas Red dextran (Thermo Scientific) in PBS was added to one well of the device and allowed to flow across the collagen within the device. Dextran flow was imaged at the device center with 5x/0.16 objective on a Zeiss Axio Observer Z1 widefield microscope every 10 minutes for 24 hours, then analyzed by generating a kymograph with custom MATLAB (MATLAB 2021b; Mathworks, Natick, MA) code. A line of best fit was generated for each binarized kymograph (threshold = 0.5) and the slope, representing the interstitial flow rate, was calculated. Slopes across dual hydrogel and single hydrogel devices were compared to assess differences in flow rate.

### Fluid flow analysis

To determine the perfusion flow profiles within acellular and epithelialized microchannels, fluorescently labeled polystyrene microspheres (500 nm Dragon Green, Bangs Laboratories, Inc) were perfused through the devices at 5 µL/min. The microchannels were imaged with a Dragonfly spinning disk confocal microscope (Andor) using the finite burst imaging protocol to achieve up to 400 frames per second using a 10x/0.45 objective. Particle tracking was performed with the Trackmate Particle Image Velocimetry (PIV) plug-in on ImageJ to calculate microsphere velocities.

### Statistical analysis

For interstitial flow with collagen hydrogels and diameter measurements, Student’s t-tests were performed. Interstitial flow with fibrin hydrogels and sprout frequency comparisons were performed using Welch’s t-test due to unequal variance between test groups. All statistical analysis was done using an alpha value of 0.05 in Prism 9 (GraphPad Software).

## Results

### Straightforward fabrication of hydrogel zones along a continuous microchannel

Fabrication of PDMS microfluidic devices often requires specialized equipment and facilities, which hinders translation to non-expert labs. We aimed to create a device that could achieve complex zonal patterning while only requiring standard benchtop equipment. To that end, we used 3D printed device molds with added hypodermic tube to cast PDMS devices, avoiding the need for photolithography techniques and equipment (**Figure 1**A). The 3D printed piece served to mold the void space for the hydrogel, while the hypodermic tube created guide channels within the PDMS to align an acupuncture needle for molding a microchannel within the hydrogel. These guide channels also served to connect the media reservoirs to the hydrogel stroma and microchannel. Hydrogel prepolymer solutions were added through the injection ports and polymerized around the acupuncture needle to generate a single microchannel running the length of the device when the needle was removed. Microchannels of at least 15 mm long (**Figure 1**B) could be robustly fabricated with multiple distinct hydrogel zones using this orthogonal injection design. The 3D-printed molds were designed with multiple hydrogel injection ports to enable the patterning of different stromal regions along the length of the microchannel, which has not been shown before. By slowly injecting hydrogel prepolymer solutions serially through each injection port, discrete hydrogel regions were formed with minimal mixing at the interface of the two gels, as visualized with fluorescent microspheres (**Figure 1**C). To ensure the hydrogels would anneal but maintain their own properties, the prepolymer solutions were injected prior to complete polymerization of the neighboring region, about 10-30 seconds apart (15). The air relief ports built into the system allowed air to escape from between the hydrogels during injection, thereby preventing interface disruption from air bubbles or gel mixture by providing a low-resistance path for the air and hydrogel solution to flow.

**Fig. 1.**
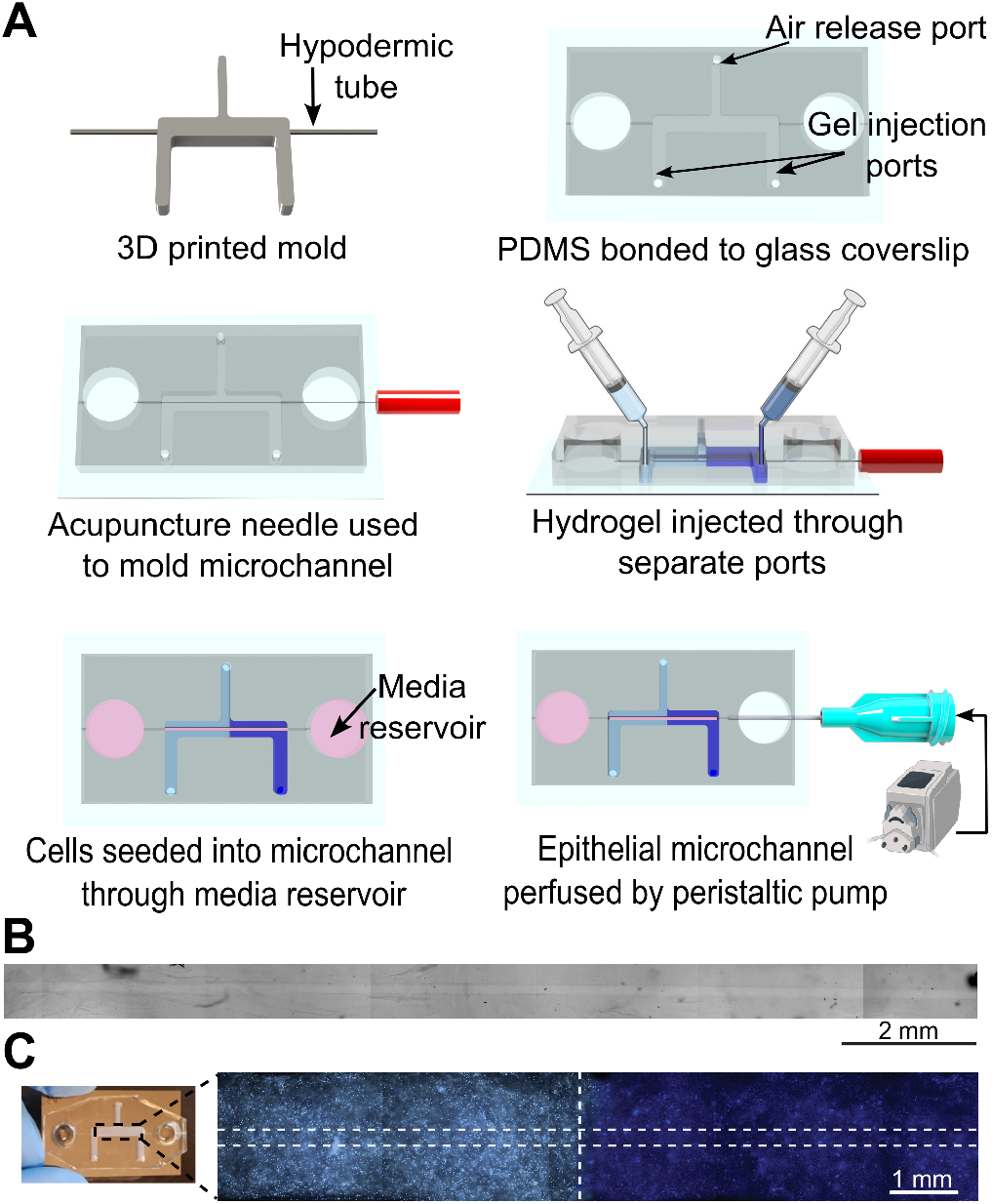
Fabrication and perfusion of a continuous central microchannel within a regionally patterned microchannel device. (A) Fabrication process for device. (B) Brightfield image of a hydrogel microchannel within a device before addition of cells (tiling artifacts can be noted along the image). (C) Fluorescent labeling with microspheres of patterned collagen hydrogel device. Horizontal dashed lines indicate edges of microchannel. Vertical dashed line indicated interface between the two regions.

### Multigel stromal regions successfully annealed together

Whereas previous work has shown that two independently injected collagen-I hydrogels anneal together to permit cellular motility between the two (15–17), we wanted to confirm small molecule flow would also be permitted across the hydrogel interface. We tracked the flow front of 70 kDa Texas Red dextran through the hydrogel in devices without a microchannel and measured the flow rate across the interface (**Figure 2**A). Devices with two patterned 3 mg/mL collagen hydrogel regions (dual) were compared to control devices with one 3 mg/mL collagen hydrogel region (single). Timelapse imaging (**Figure 2**B) was used to capture the flow front and kymographs were generated to visualize Texas Red dextran flow rates across the devices (**Figure 2**C). We quantified flow rates from the kymograph slopes and found no significant difference in flow rates between the dual hydrogel (5.316±2.46 µm/min) and single hydrogel (4.955±1.413 µm/min) devices (**Figure 2**D). This indicated the collagen gels successfully annealed and the interface between the gels did not cause disruptions in interstitial flow.

**Fig. 2.**
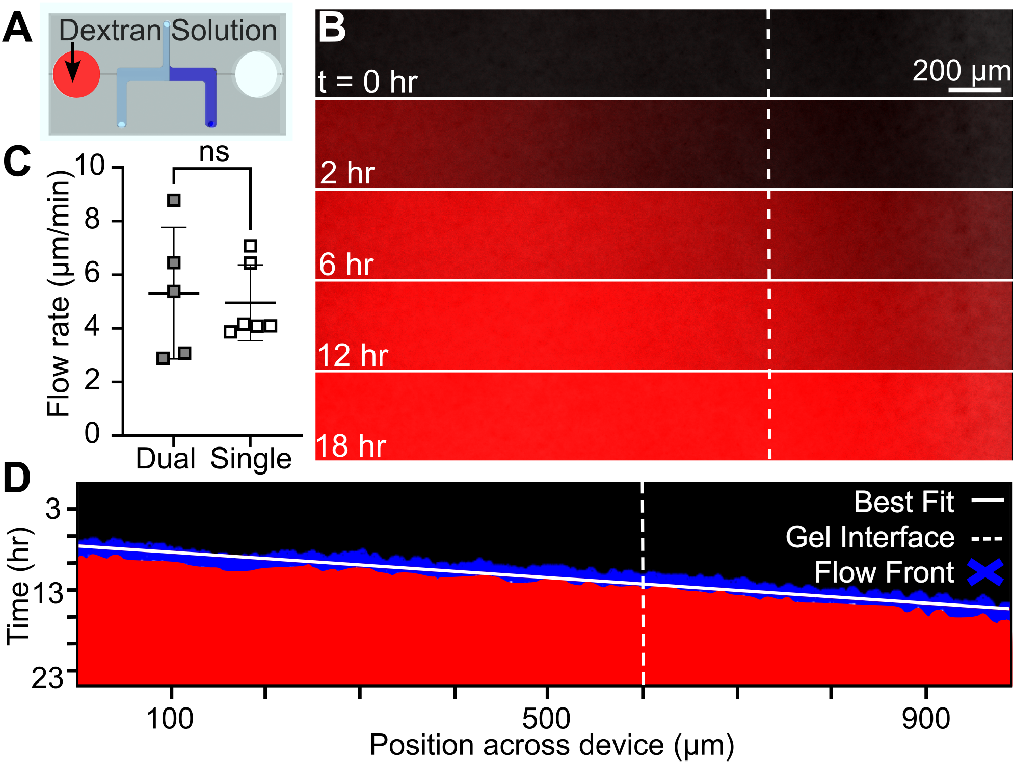
Interface of collagen gels does not impact diffusion through the gel. (A) Schematic of Texas Red dextran interstitial flow assay set-up. (B) Fluorescent images of Texas Red dextran diffusion front at various time points. Vertical dashed line indicates interface between collagen hydrogels. (C) Comparison of Texas Red dextran diffusivity in single dual hydrogel devices. (ns = no significant difference; n ≥ 5; p = 0.776, α = 0.05). (D) Binarized kymograph of Texas Red dextran diffusion front with line of best fit for the position of the front over time.

### Microchannel structure is unaffected by the gel interface

When removing the acupuncture needle, forces are applied to the surrounding hydrogel which could potentially cause de-lamination of the hydrogel interface or imperfections in the microchannel molding. Single and dual hydrogel region devices were made to evaluate if patterned hydrogel devices were affected by the demolding process. No delamination or distortion was evident at the interface in patterned devices (**Figure 3**A). Microchannel diameters were measured along the device length and normalized to the acupuncture needle diameter and no significant difference was found between single and dual hydrogel devices (**Figure 3**B). The average difference in diameter between the channel and acupuncture needle along the length of each microchannel was 9.5 µm and 5.4 µm for single and dual hydrogel devices, respectively (**Figure 3**C). While there were no apparent visual or dimensional defects to the molded microchannels, we also quantified fluid flow profiles through the microchannel, particularly across the gel-gel interface. To visualize the fluid flow, dual hydrogel devices were perfused with 500 nm fluorescent microspheres, which served as Lagrangian fluid tracers. Timelapse images of microsphere flow through the device were taken at the hydrogel interface, and subsequent PIV analysis revealed the velocity of each microsphere as a function of its position in the channel (**Figure 3**D). Devices were found to have a parabolic flow profile with no disruptions at hydrogel zonal interfaces, indicative of unobstructed laminar flow (**Figure 3**E).

**Fig. 3.**
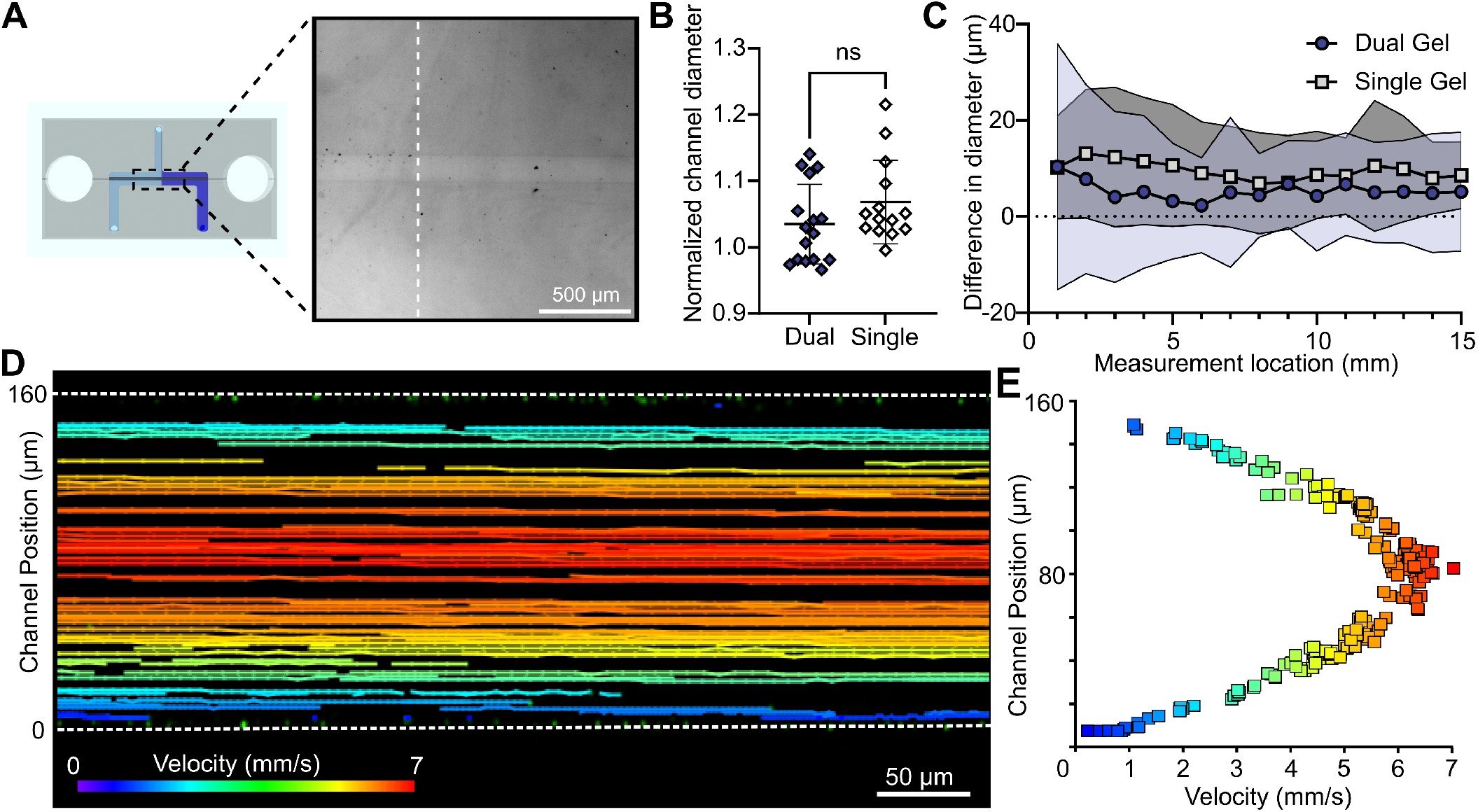
Collagen interface does not impact the microchannel structure or laminar flow profile through the interface. (A) Microchannel structure at the hydrogel interface within a dual hydrogel device. (B) Comparison of mean normalized channel diameters between single and dual hydrogel devices. (n≥s = no significant difference; n 14; p = 0.148; α = 0.05) (C) The mean difference between the diameter of the microchannel and acupuncture needle at each measurement location of single and dual hydrogel devices. Shaded region indicates SD. (D) PIV tracks of microsphere flow within a dual hydrogel device microchannel. Dashed lines indicate channel boundaries. (E) Fluid flow profile in a microchannel in a dual hydrogel device.

### Epithelialization of the microchannel is unaffected by hydrogel patterning

We next sought to epithelialize the microchannel within the patterned devices. Following demolding of the microchannel, MDCK cells were introduced via a hydrostatic pressure gradient between the two media wells. To ensure robust epithelialization of the microchannel, cell-laden devices were cultured under static conditions for five days and then subjected to 24 hours under physiologic flow conditions (**Figure 4**A). After culture, the microchannels were fully cellularized with a complete monolayer. Immunofluorescent staining revealed junctional E-Cadherin and apical F-Actin, which were indicative of epithelial polarization (18) (**Figure 4**B). Additionally, no cells migrated along the gel interface perpendicular to the microchannel, further supporting that the patterned gels were fully annealed together. Finally, we assessed the fluid flow through the established epithelial tube with perfusion of 500 nm diameter fluorescent microspheres. PIV analysis revealed that flow in the epithelialized microchannel was laminar with a characteristic parabolic flow profile (**Figure 4**CD).

**Fig. 4.**
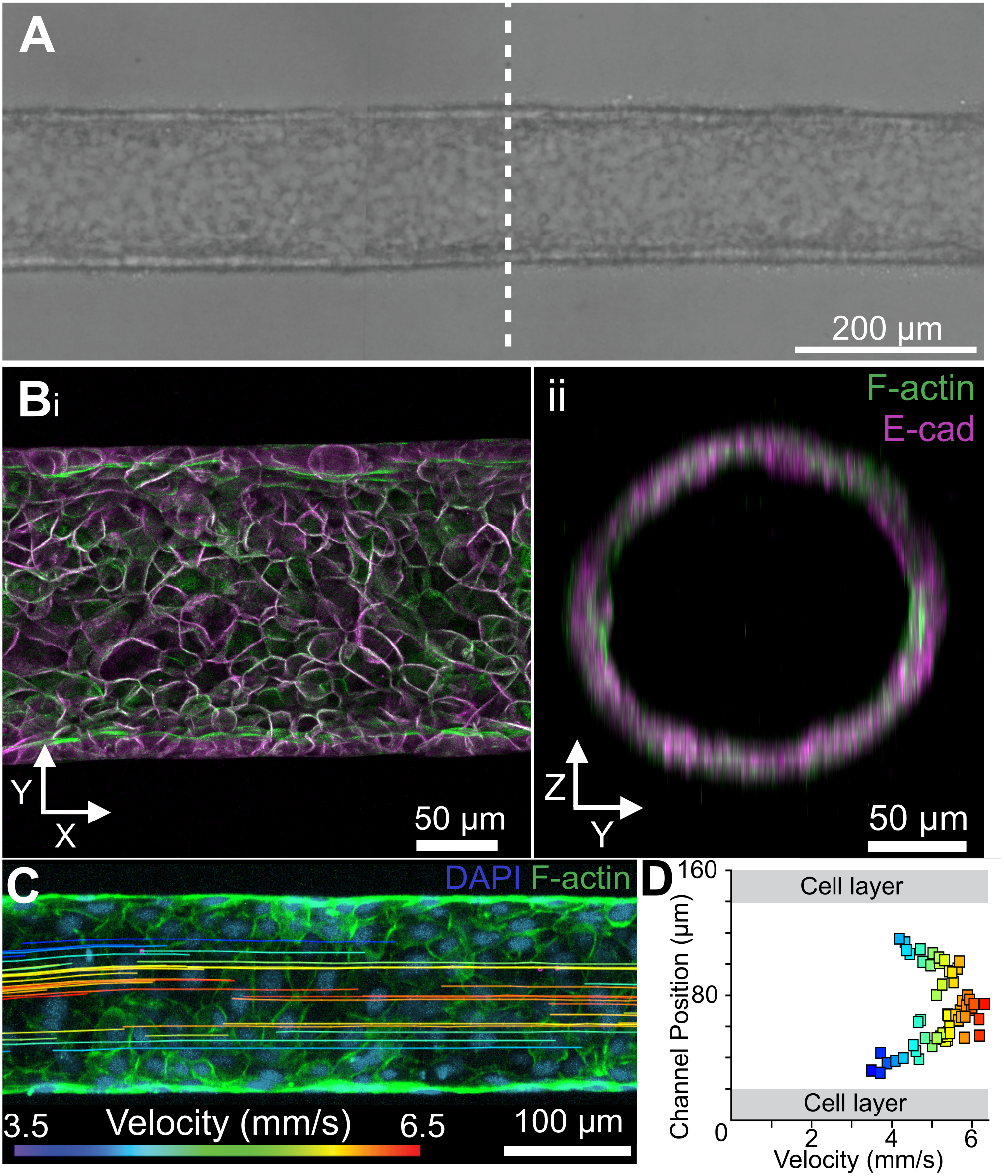
Hydrogel interface does not disrupt epithelialization of microchannel. (A) Epithelialized microchannel within a patterned device. Dashed line indicates hydrogel interface. (B) Maximum projection of epithelialized microchannel within a patterned device in the (i) (X,Y) plane and the (ii) (Y,Z) plane.(C) Microsphere flow profile within a dual hydrogel patterned device with an epithelialized microchannel. (D) PIV microsphere pathlines within dual hydrogel patterned device with an epithelialized microchannel.

### Multiple sequential regions and hydrogel compositions can be patterned along a central epithelial tube

The primary goals of this fabrication methodology were to create a device that was easy to construct and usable to investigate a variety of questions about local cell-cell and cell-ECM interactions. We, therefore, tested this system’s compatibility with different common hydrogel compositions and the potential for increased zonal patterning complexity. To assess these capabilities, we made devices with fibrin hydrogels and with collagen hydrogel zones of differing concentrations. Dual fibrin hydrogel devices were made via the same methods as previous collagen devices, wherein 5 mg/mL fibrin hydrogels with encapsulated fluorescent microspheres were injected serially into the device to create a regionally patterned fibrin device (**Figure 5**A). Patterned fibrin devices without microchannels were used to assess small molecule interstitial flow across the interface and compared to single-gel fibrin devices, as done previously with collagen devices. There was no significant difference in interstitial flow rate between the dual (4.85 ± 0.71 µm/min) and single (5.84 ± 2.56 µm/min) fibrin hydrogel devices (**Figure 5**B). Dual fibrin devices with microchannels were compared with single gel devices to validate that patterned fibrin gels did not delaminate or distort during microchannel demolding (**Figure 5**C). As all previous experiments focused on assessing the gel patterning and microchannel molding technique within uniform hydrogels, we also wanted to ensure that hydrogels with differing properties could be used in this system as this would enable future investigation into the effects that variable stromal biophysical and biochemical properties have on developmental patterning. We used 3 mg/mL and 6 mg/mL collagen to pattern zones within the devices (**Figure 5**D), and found no significant difference in molded microchannel diameters between the different collagen concentration regions of the same device (**Figure 5**E). To assess the ease with which these methods could be adapted to increase the number of zones along the epithelial microchannel, we designed a 3D-printed mold with two additional hydrogel injection ports and air outlets. We successfully created devices with four patterned hydrogel regions with minimal modification to the fabrication methodology (**Figure 5**Fi/ii).

**Fig. 5.**
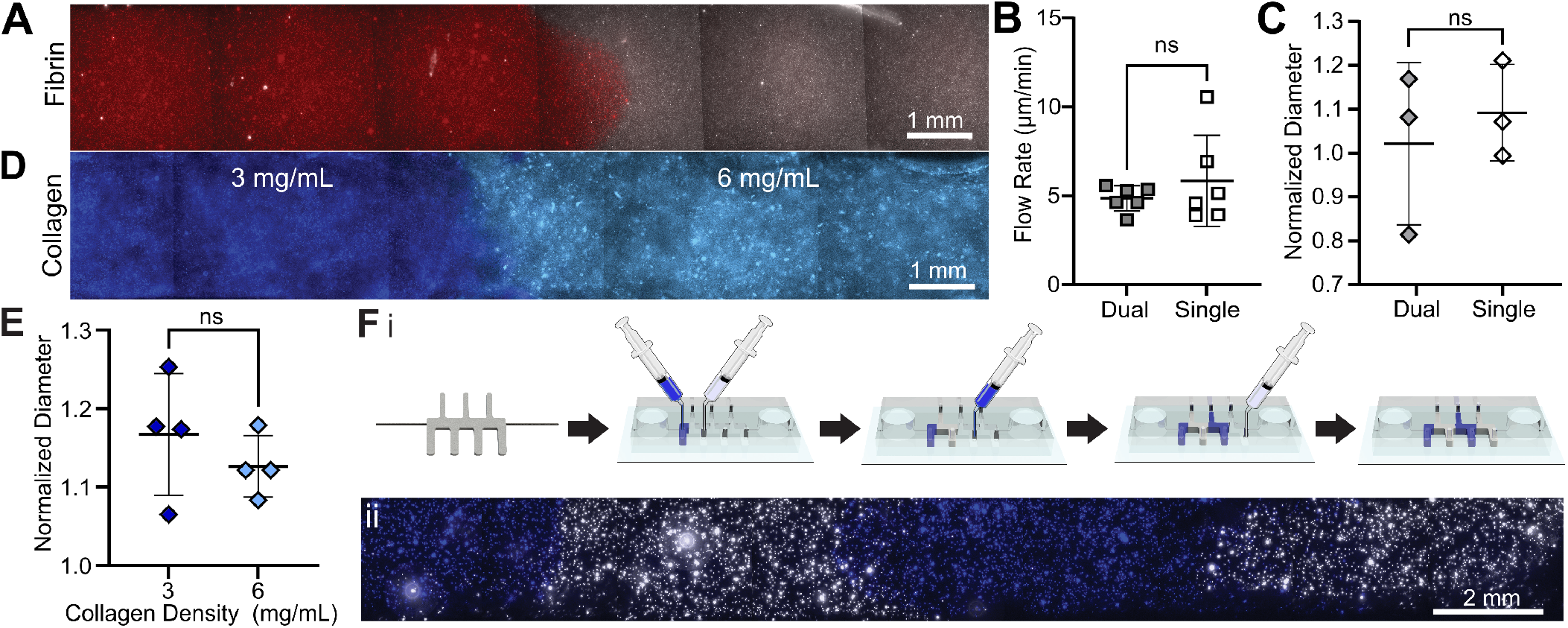
Device design is compatible with various hydrogel formulations and can be adapted to increase the number of hydrogel regions. (A) Dual patterned fibrin hydrogel device with distinct fluorescent microspheres in each region. Comparison of (B) fluorescent dextran diffusion (ns = no significant difference; n = 6; p = 0.398; α= 0.05) and of (C) microchannel diameter in in dual and single fibrin hydrogel devices (ns = no significant difference; n = 3; p = 0.599; α= 0.05). (D) Dual patterned multi-density collagen hydrogel device seeded with distinct microspheres to demarcate the two density regions. (E) Comparison of microchannel diameter in the two collagen regions (ns = no significant difference; n = 4; p = 0.384; α= 0.05). (F) i. 3D printed mold design and device fabrication for patterning device with four collagen hydrogel regions. ii. Fluorescent image of four region patterned device visualized with encapsulated fluorescent microspheres in each region.

### Epithelial phenotype is locally altered by zonal stromal cell patterning

In addition to variable zonal hydrogel composition, stromal cell populations can be regionally patterned along the microchannel to investigate regional cell-cell interactions. 3T3 fibroblasts were chosen for this investigation of local cellsignaling as they have been shown to induce tubulogenic sprouting in MDCKs when cultured together (19, 20). Two region devices were patterned with a 3T3 fibroblast-laden collagen hydrogel and an acellular collagen hydrogel along a microchannel seeded with MDCKs (**Figure 6**A). 3T3s were viable within the hydrogel region over the entire culture period and took on a spread morphology within 2-3 days of culture (**Figure S1**). MDCKs epithelialized the microchannel to form a continuous monolayer (**Figure 6**B) within 5-7 days, similarly as when seeded without stromal cells (**Figure 4**). Following complete epithelialization of the microchannel, MDCK sprouts were observed to invade, or ‘sprout’ into the stromal hydrogel area of the device (**Figure 6**Ci/ii). To quantify the epithelial response to regional fibroblast patterning, sprouts were counted across the entire microchannel after 7 days of static culture. The distribution of sprouts from all devices shows an overall increase of sprouts in from the epithelium in the cellular stromal region (**Figure 6**D) compared to the acellular region. There was a significantly greater average number of epithelial sprouts within the cellular stromal zone (14 sprouts) compared to the acellular stromal zone (6 sprouts) of each co-culture device (**Figure 6**E).

**Fig. 6.**
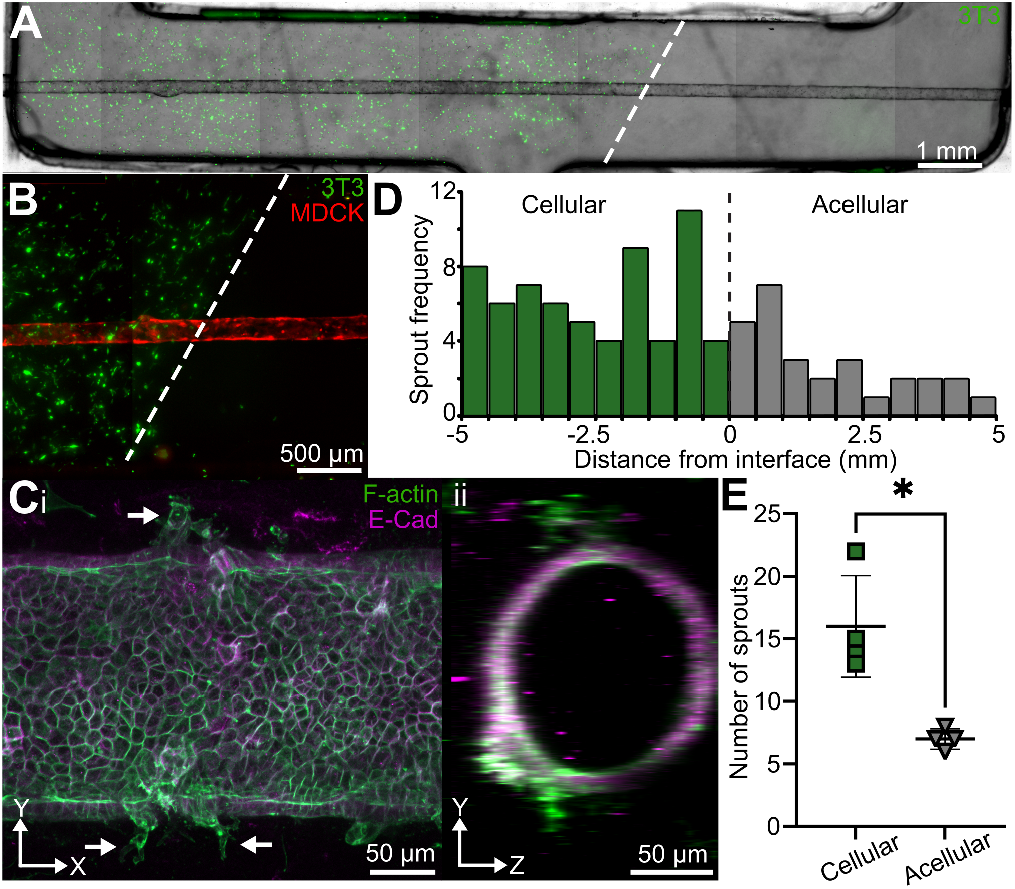
Device supports cell-cell signaling between stromal populations and the epithelium. (A) Fluorescent and brightfield overlay of 3T3 patterned device with an MDCK epithelialized microchannel. Dashed line indicates hydrogel interface.(B) GFP expressing 3T3 patterned device at the interface between the two regions (dashed line). (C) Maximum projection of sprouted (arrows) epithelialized channel within the 3T3 stromal region in the (i) (X,Y) and (ii) (Y,Z) planes. (D) Total frequency and location of all MDCK sprouts along epithelialized microchannels. (E) Average MDCK sprout number in 3T3 region versus acellular region. (*p = 0.0195; n = 4; α = 0.05)

## Discussion

The architecture of many major organs is defined by highly branched tubular epithelium, including the kidney, lungs, and pancreas, which arise through the process of branching morphogenesis during development. The epithelial tubes within these developing tissues are further characterized by regionally distinct cell populations. In the kidney, this is exemplified by the nephron’s many segments and 17 unique cellular populations; similarly, the lung is characterized by zones of cell types patterned along the upper to lower respiratory tract. Although the developmental stages and cellular functions within tubular organs are well characterized, much remains unknown about the factors regulating differentiation and specification events spatially along these tubular epithelial tissues. It is thought that the mechanical cues from the ECM and biochemical cues from neighboring cells play a large role in organogenesis (21–23). Indeed, proximodistal patterning of the lung epithelium is known to be influenced by localized differences in the microenvironment, including mesenchymal cell identity and mechanical forces (24–27). Additionally, microenvironmental changes associated with the spatially constrained expression of ECM modifying enzyme lysyl oxidase (LOX) along the cortical-medullary axis of the kidney may be involved in nephron segment specification (8). These developmental processes are inherently diffi-cult to study and there is limited control of the many variables that exist within *in vivo* and *ex vivo* systems. *In vitro* systems allow tight control over multiple variables; however, the existing *in vitro* systems do not facilitate the study of regional microenvironmental influences. The methods herein address this gap by enabling zonal patterning of stromal regions along a cellular microchannel to tease apart how stromal cells and extracellular matrix properties influence epithelial phenotype along a continuous epithelial tube. We show that this MPS is adaptable to multiple hydrogel systems and number of stromal regions (**Figure 5**) with minimal alterations in the patterning methodology. Importantly, we also show the ability to spatially control cell-cell interactions between a stromal and epithelial cell population to produce regional phenotypic changes within our model (**Figure 6**). While our focus herein has been on creating tubular epithelial tissue, this system could similarly be used to probe questions regarding vascular function and the endothelium, such as tumor-induced endothelial cell changes or factors influencing organ-specific endothelial phenotypes (28, 29).

Layer-by-layer fabrication is a popular technique to create constructs with vertically or horizontally stacked hydrogel layers mimicking stratified solid tissues (30–34). Sequential addition of hydrogels with unique prepolymer and crosslinker concentrations are used to create composite hydrogels with discrete regions of varying biochemical compositions and mechanical properties (30, 35, 36). Yet, microfluidic models for epithelial or endothelial tissues have not incorporated these techniques to create tissue heterogeneity along microchannels. We thus adapted the layer-by-layer technique to create horizontal hydrogel layers, or regions, with the added complexity of a central cellular microchannel. Furthermore, many layered hydrogel composites are made via 3D bioprinting (32, 37–41). These methods can be both costly and difficult for non-experts to use and often lack the resolution needed to create microchannels within bulk hydrogels. Other microfluidic systems similarly rely upon complex microfabrication and photolithography techniques to generate devices and channels with the structure and resolution desired (42–46). However, the approach described herein does not require 3D bioprinting techniques or cleanroom microfabrication experience. Instead, by molding microchannels with commonly available supplies and patterning regions through manual injection and timed polymerizations, our design methodology is easily transferrable and accessible to biological laboratories with limited engineering or technical background in microfabrication. Furthermore, the equal interstitial flow rates of fluorescent 70 kDa dextran through dual and single hydrogel region devices (**Figure 2**) and low variability of microchannel diameter across the length of the device (**Figure 3**) demonstrate that our simple, inexpensive methodology is robust and capable of producing a patterned stroma along a perfusable microchannel.

Due to the critical role the biophysical and biochemical properties of the ECM play in orchestrating cellular function and tissue patterning (47–51), careful consideration is required when choosing an appropriate hydrogel system to recapitulate the ECM of a tissue of interest. Therefore, we validated our system with collagen hydrogels of multiple densities as well as fibrin hydrogels, as these natural protein-based hydrogels are frequently used within other MPSs due to high cytocompatibility and the ability for remodeling by the cells within the gels (52, 53). The methodology to create multiple hydrogel regions proved equally compatible with fibrin hydrogels and collagen hydrogel of differing densities, which suggests the use of other hydrogels of interest, including synthetic or photopolymerizable gels, could be similarly successful. The addition of photopolymerizable gels is especially interesting as the photochemistry allows precise spatial control of hydrogel properties. This would be useful in creating hydrogels with high spatial resolution gradients of mechanical properties and bioactive ligands (54–57) which can be used to mechanistically understand tubular organogenesis. The forces generated by fluid flow through the developing and mature tubular organs are known regulators of epithelial function (58). For example, fluid flow within the renal epithelium regulates expression of multiple transporter channels such as Na/K-ATPase and aquaporin-1 and −2 to control and maintain the ionic homeostasis of the blood (59, 60), which is central to the nephron’s function. Given the importance of fluid flow, we validated that the hydrogel interface within our model did not interfere with perfusion in both acellular (**Figure 3**) and epithelialized (**Figure 4**) microchannels. Complete epithelialization of the microchannel was successful and remained intact under flow rates with the applied shear stress of 1-2 dyne/cm2, equivalent to the shear rate in kidney tubules, venous vessels, and the developing lung (61–64). Phenotypic apical-basal polarization of F-actin and Ecadherin expression was observed with no abnormalities in lumen structure at the interface between gel regions. Moreover, the fluid flow within both the acellular and epithelialized microchannels remained laminar with a parabolic flow profile as determined by PIV analysis.

The creation of biochemical signaling gradients along the axis of a cellularized microchannel is a key strength of this model. We leveraged spatial patterning of stromal populations to create soluble factor gradients within devices of MDCKs and 3T3 fibroblasts co-cultures (**Figure 6**). The combination of MDCKs and 3T3s was chosen as fibroblasts secret hepatocyte growth factor, which is known to induce tubulogenic sprouting in MDCKs (19, 20). Within the same device, regions containing 3T3 fibroblasts had more than two times the number of MDCK sprouts than in regions without a stromal cell population. These data confirmed that tubulogenic sprouting was regionally dependent and that the device design enables spatial control of biochemical cell-cell interactions. Further spatial patterning of biochemical signaling factors could be achieved within this model using laminar flow patterning by interstitial perfusion of signaling molecules into each individual stromal region through the hydrogel injection ports (65–67). Coupling patterned interstitial flow of soluble molecules with advanced synthetic hydrogel systems capable of bioorthogonal crosslinking (68) would enable *in situ* post-polymerization modification and patterning of the hydrogel’s biochemical or biophysical properties.

## Conclusion

We have developed and validated a simple strategy to create an *in vitro* microphysiological system with regionally patterned stromal cell populations and hydrogel properties along the length of a perfused continuous tubular epithelium. This highly customizable system has wide utility for modeling epithelial and endothelial tissue interactions with heterogeneous hydrogel compositions and/or stromal cell populations during development, homeostasis, or disease. This model system better recapitulates the heterogenous tissue structure and cellular organization of *in vivo* organs and can enable the dissection of specific microenvironmental factors independently, which is challenging in animal models. The potential insights into critical cell-ECM and cell-cell interactions that govern tissue patterning gained with this model may provide new strategies for regenerative medicine and biofabrication approaches to achieve functional, clinically relevant tissues.

## ACKNOWLEDGEMENTS

The authors thank Katherine M. Nelson, Ph.D., for reviewing and commenting on the manuscript. This work was supported in part by grants from the National Institutes of Health: R01DE029655 and R01HL145147. Microscopy equipment was acquired with a shared instrumentation grant (S10 OD030321) and access was supported by NIH-NIGMS (P20 GM103446, P20 GM139760) and the State of Delaware. The BioRxiv template was adapted from the Henriques lab.

## DATA AND RESOURCE AVAILABILITY

Downloadable engineering drawings and .STL files for the device 3D printed components can be found at: https://www.gleghornlab.com/resources

## AUTHOR CONTRIBUTIONS

Conceptualization (BC, BJF, JPG), Methodology (BC, BJF, JPG), Investigation (BC, BJF), Validation (BC, BJF, JPG), Visualization (BC, BJF, JPG), Formal Analysis (BC, BJF, JPG), Writing – Original Draft (BC, BJF), Writing – Review & Editing (BC, BJF, JPG), Supervision (JPG), Project Administration (JPG), Funding acquisition (JPG)

## Supplementary Information

**Fig. S1.**
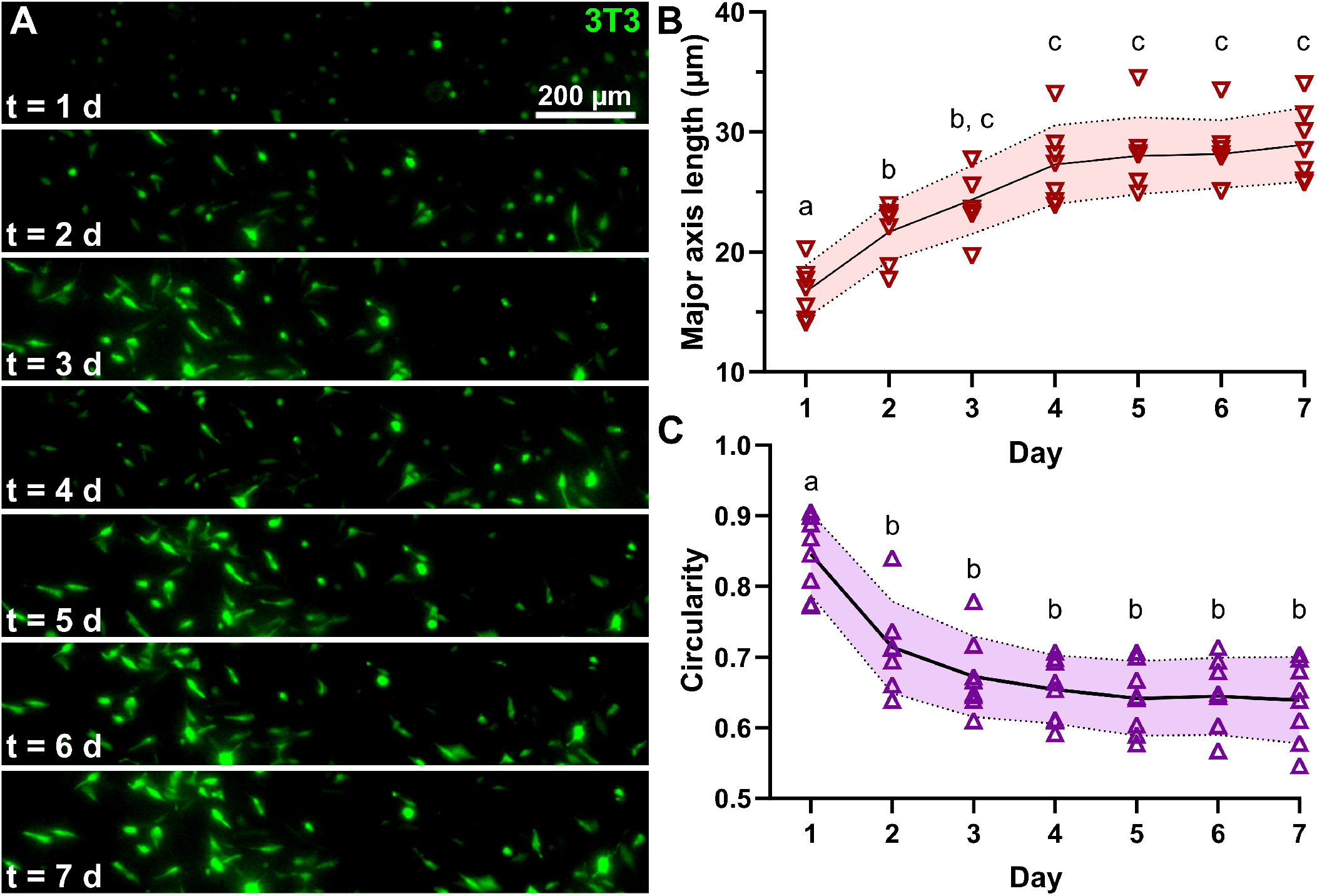
The co-cultured 3T3 fibroblasts embedded within the collagen stroma have a phenotypic spread out morphology within 2-3 days of culture. (A) Representative fluorescence images of GFP expressing 3T3 morhpology at each day of 7 day co-culture. (B) Average length of 3T3 cell spreading increases over the first 2-3 days of culture. (Days with different letters are significantly different, shaded region represents the standard deviation, ANOVA p < 0.001, n = 7, α = 0.05) (C) The average circularity of the 3T3 cells decreased significantly after the first day. (Days with different letters are significantly different, shaded region represents the standard deviation, ANOVA p < 0.001, n = 7, α = 0.05)

**Fig. S2.**
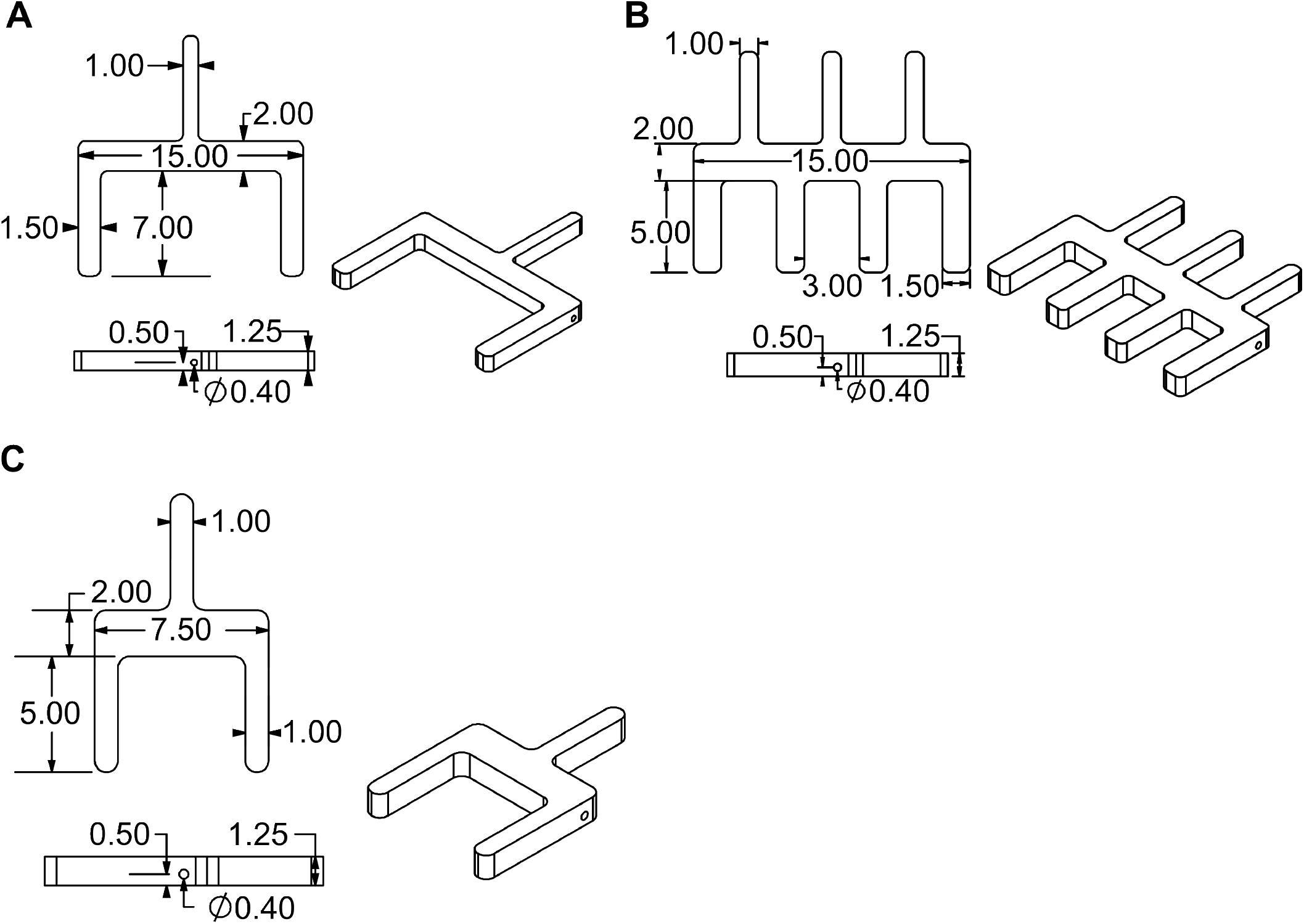
Engineering drawing of 3D models used to mold devices in PDMS. All dimensions are in millimeters (A) Design for 2 region large device. (B) Design for 2 region short device. (C) Design for 4 region device. Downloadable STL files can be found at: https://www.gleghornlab.com/resources

## Notes

### Competing Interest Statement

The authors have declared no competing interest.

